# InsectBrainDatabase - A unified platform to manage, share, and archive morphological and functional data

**DOI:** 10.1101/2020.11.30.397489

**Authors:** Stanley Heinze, Basil el Jundi, Bente G. Berg, Uwe Homberg, Randolf Menzel, Keram Pfeiffer, Marie Dacke, Eric Warrant, Gerit Pfuhl, Jürgen Rybak, Kevin Tedore

## Abstract

Insect neuroscience generates vast amounts of highly diverse data, of which only a small fraction are findable, accessible and reusable, despite open data mandates by funding bodies. We have therefore developed the InsectBrainDatabase (*IBdb*), an open platform for depositing, sharing and managing a wide range of insect neuroanatomical and functional data. It facilities biological insight by enabling effective cross-species comparisons, by intimately linking data on structure and function, and by serving as hub for information on insect neuroethology. The *IBdb* provides novel visualization and search tools, which are also available in a unique private mode of the database, before data is made public. This allows users to manage and visualize unpublished data, creating a strong incentive for data contribution and eliminating additional effort when publicly depositing the data at a later stage. These design principles could also serve as a blueprint for similar databases in other fields.

## Introduction

Data are the essence of what science delivers - to society, to researchers, to engineers, to entrepreneurs. These data enable progress, as they provide the basis on which new experiments are designed, new machines are developed, and from which new ideas emerge. Independent of the research field, many terabytes of data are produced every year, yet only a small fraction of these data become openly available to other researchers, with even less penetrating the invisible wall between the scientific community and the public (***Mayernik, 2017***). While research papers report conclusions that are based on data and present summaries and analyses, the underlying data most often remain unavailable, despite their value beyond the original context. Whereas this is changing in many fields and the use of open data repositories becomes increasingly mandatory upon publication of a research paper, this is not ubiquitous and older data remain inaccessible in most instances. Additionally, merely meeting the data deposition requirement by ‘dumping’ poorly annotated raw files on an internet platform does not aid transparency or reuse of the data. To ensure common standards for data repositories and the datasets to be stored in them, the FAIR principles for data deposition (Findability, Accessibility, Interoperability, and Reusability) were developed (***Wilkinson et al., 2016***). It is clear from these principles that annotation and rich meta-data are essential, if a dataset is supposed to be beneficial to others. While this is relatively easily achievable for data such as gene sequences, protein sequences, or numerical datasets, the challenges are much bigger for complex morphological data, physiological observations, or behavioral studies. The difficulties result not only from large file sizes of image stacks, high-speed videos, or recorded voltage traces, but also from the heterogeneous data structure often generated by custom designed software or equipment.

Insect neuroscience is no stranger to these challenges. Particularly for research outside the genetically accessible fruit fly *Drosophila*, no universal data repository exists that allows retrieval of original observations that underlie published articles. Research groups worldwide investigate the nervous systems of a wide range of insect species, but mostly operate in isolation of each other. Data from these projects are often deposited in local backup facilities of individual institutions and thus remain inaccessible to the community. Combined with a lack of interoperability caused by independent choices of data formats this has the potential to severely hamper progress, given that interspecies comparison is one of the existential pillars on which insect neuroscience rests. The problem is amplified by the fact that much of the data are both large and heterogeneous (e.g. 2D and 3D images, models of brain regions, digital neuron reconstructions, immunostaining patterns, electrophysiological recordings, functional imaging data).

A final problem is that depositing well-annotated data takes time and effort, and little incentive is generally given to prioritize this work over acquiring new data or publishing research papers. While this is true for all research fields, the complex data in neuroscience requires an extra amount of effort to meet acceptable standards. This has made depositing data in a form that is useful to the community a relatively rare event. Early efforts were made to develop brain databases for various insect species (e.g. honeybee (***Brandt et al., 2005***), Manduca sexta (***el Jundi et al., 2009b***), Tribolium castaneum (***Dreyer et al., 2010***), desert locust (***Kurylas et al., 2008***)), but in those cases, the anticipated interactive platforms for exchange and deposition of anatomical data were short-lived and not used beyond the laboratories that hosted them. More recently, several successful databases for anatomical data from insects were developed. Most notably, Virtual Fly Brain (VFB) (***Osumi-Sutherland et al., 2014***) now bundles most efforts in the *Drosophila* community regarding the deposition of neuroanatomical data - including single cell morphologies from light microscopy, data from recent connectomics projects (most notably from ***Scheffer et al. (2020)***), as well as catalogues of GAL4 driver lines, which enable access to specific neurons with genetic methods. VFB hosts most content of older independent databases, such as FlyCircuit (***Chiang et al., 2011***) and FlyBrain (***Armstrong et al., 1995***) and is dedicated to providing smart ways of utilizing and visualizing *Drosophila* neuroanatomy. Similarly, but more focused on data visualization and connectivity modeling, theFruitFlyBrainObservatory allows access to current datasets of single neuron morphologies from *Drosophila*. Another database, founded in 2007, has grown substantially over recent years: NeuroMorpho.Org (***Ascoli et al., 2007***). It provides 3D reconstructed datasets of more than 100,000 neurons from across animal species and includes substantial numbers of single cell data from insects. While the latter platform is comparative in nature, it does not offer dedicated tools for comparisons between species or much context for the deposited neuron skeletons. In contrast, the data on VFB are much richer and different datasets are tightly linked to each other, allowing, for example, correlation between GAL4 driver lines and electron microscopy based single neuron reconstructions. Yet, no comparison to other species is possible or intended via VFB. Additionally, no systematic information is provided about the function of the deposited neurons in either database, precluding insights into the structure-function relations that are critically important to understanding the insect brain.

To address these shortcomings we have developed the InsectBrainDatabase (*IBdb*), a cross-species, web-based platform designed for depositing and sharing research data that consist of morphological and functional information from insect brains. With an overall modular design, a concept for dual use as depository and data management tool, combined with widely useful visualization tools, this database yields a tool for increasing transparency, accessibility and interoperability of insect neuroscience data. Moreover, the newly developed concepts are not only relevant to insect neuroscience, but to any scientific field that can be linked to a hierarchically organized, ontological framework. We thus hope that our conceptual design can be adopted by a range of users from across the sciences to simplify data handling and make scientific results in general more transparent.

## Results

### Database outline

The ‘Insect Brain Database’ (*IBdb*) can be found on the internet at insectbraindb.org and is freely available to everyone. It can be used with most modern web browsers with active Javascript (tested with Google Chrome, Safari, Firefox) on computers running any operating system, without the need for any additional plugins. A user account can be registered free of charge and is required for users who wish to download data and to contribute content.

The *IBdb* is divided into three main hierarchical layers: Species, brain structures, and neuron types. Each level is additionally linked to ‘experiments’, which is the fourth major organizational layer of the database (***Figure 1***A). At each level, a database entry is represented by profile pages, on which all relevant information is collected (***Figure 1***C). These profile pages are the core of the database. They can be reached directly by several search functions, as well as by entry lists. As profile pages are embedded in a hierarchical framework of species, brain regions, and cell types, they become automatically associated with metadata. For instance, an entry for a pontine neuron of the fan-shaped body (central body upper division) in the monarch butterfly would become linked to the brain regions ‘fan-shaped body/central body upper division’ and its parent region ‘central complex’, as well as to ‘monarch butterfly’ as species. The neuron can then be found, for example, by querying the brain regions associated with it, or by exploring the species of interest.

**Figure 1.**
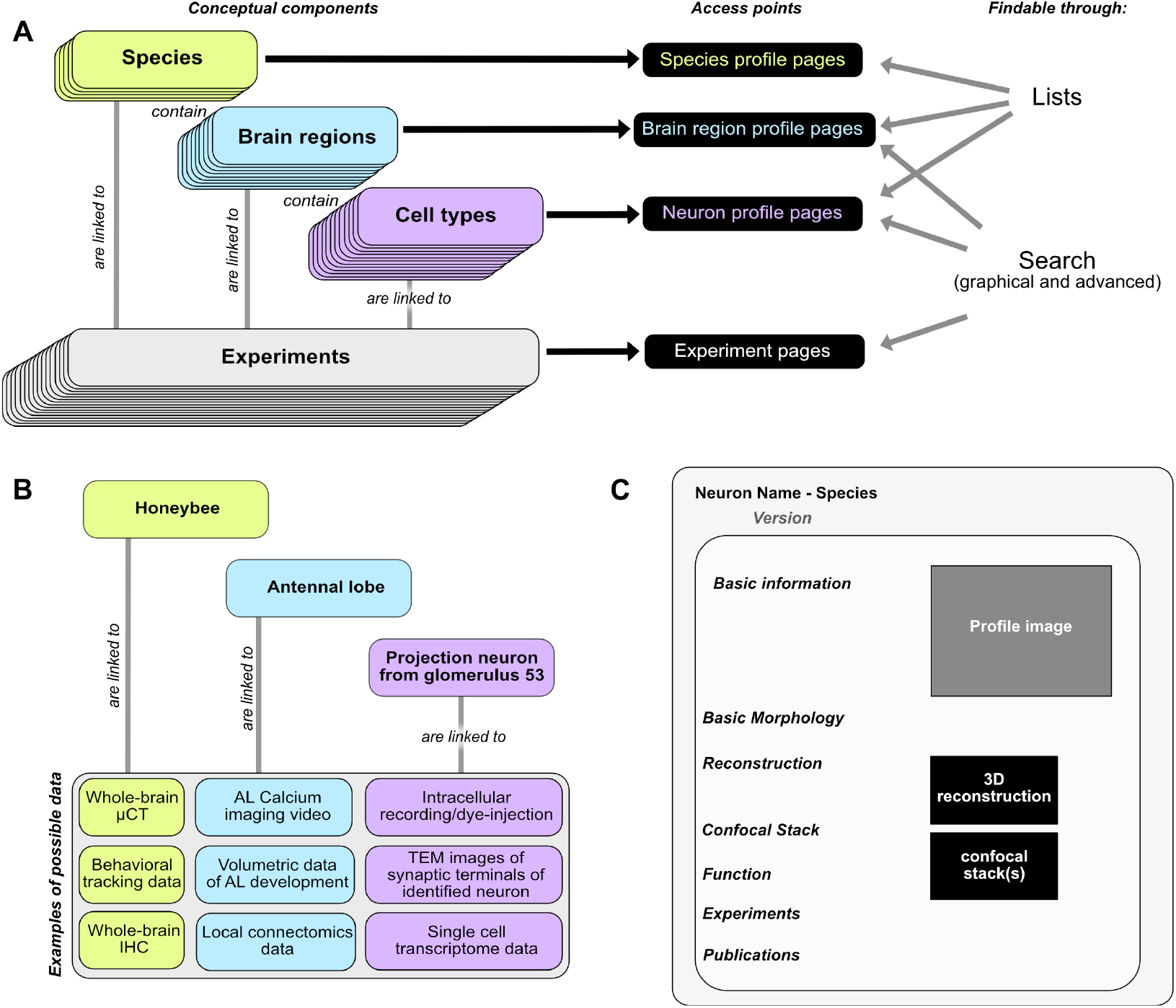
Basic concepts behind the Insect Brain Database. A. Organizational layers of the database. Elements in each layer are represented on their respective profile pages, which can be located either through lists or search interfaces. B. Examples for information that can be deposited on each level of the database, illustrating how diverse data is automatically associated with hierarchical metadata. C. Schematic illustration of a neuron profile page. Detailed anatomical and functional data is available on this page, from where experiments associated with this cell type are also linked. Similar pages exist for species and brain regions. **Figure 1-Figure supplement 1.** Technical overview of database layout.

Species entries contain representative data of an insect species, with the aim of defining that species and the overall layout of its brain. Similarly, entries for brain regions and cell types contain representative examples of data that illustrate the respective entity and provide all information to unambiguously define it, essentially providing type-specimen. Contrary to the first three levels, experiment entries contain specific data from individual, defined experiments (***Figure 1***B). As research data can be obtained at the levels of species, brain regions and cell types, experiment entries exist for all three levels and are accessible via the respective profile pages (***Figure 1***A). This distinction between representative and concrete data is important, as for example only one entry for a certain columnar neuron type of the central complex exists in the database, yet, if that single cell type was subject to 30 intracellular recordings, the profile page of that cell type would list 30 experiments, each containing a unique individual neuron with its associated physiology. The information on the profile page represents the consensus results of all experiments and can thus evolve with new experiments being added.

In the context of this evolving content, it is key to ensure that entries which are cited in published work remain findable in the exact form they existed when they were cited (***Ito, 2010***). Therefore each entry receives a persistent identifier. This identifier (a ‘handle’) links to a version of the entry that was frozen at the time it was created (and cited). Once information is added or removed from that entry, a new handle must be generated, providing a new access link for future citations. This system is applied to multiple levels of the database (experiments, neurons and species) and ensures that all information in the database, as well as the interrelations between entries, are truly persistent.

### Interactive search interfaces

As locating specific datasets is one of the core functions of a database, we have developed a novel, more intuitive way to find specific neuron data. A graphical representation of the insect brain, resembling the overall anatomical outline of all brain regions (***Figure 2**A*), makes it easy to search for neurons within single species and across species. This graphical interface is generated directly on the database website for each species and is adapted from a generic insect brain, i.e. a shared ground plan. This generic brain is the least detailed fall-back option for any cross-species search and was developed based on the insect brain nomenclature developed by ***Ito et al. (2014)***. It resembles the consensus anatomical hierarchy of all brain regions in insect brains. Within this hierarchy, the entire brain is divided into 13 super-regions, which consist of individual neuropils. The latter can be further divided into sub-regions. While all super-regions exist in all species, differences become more pronounced at lower levels of the hierarchy. The generic brain therefore largely contains super-regions as well as several highly conserved neuropils (***Figure 2-Figure Supplement 1, Figure 2-Figure Supplement 2***). As these categories are simply tags of brain region entries used to organize the database, the search interface does not differentiate between them, simplifying the user experience (***Figure 2***A).

**Figure 2.**
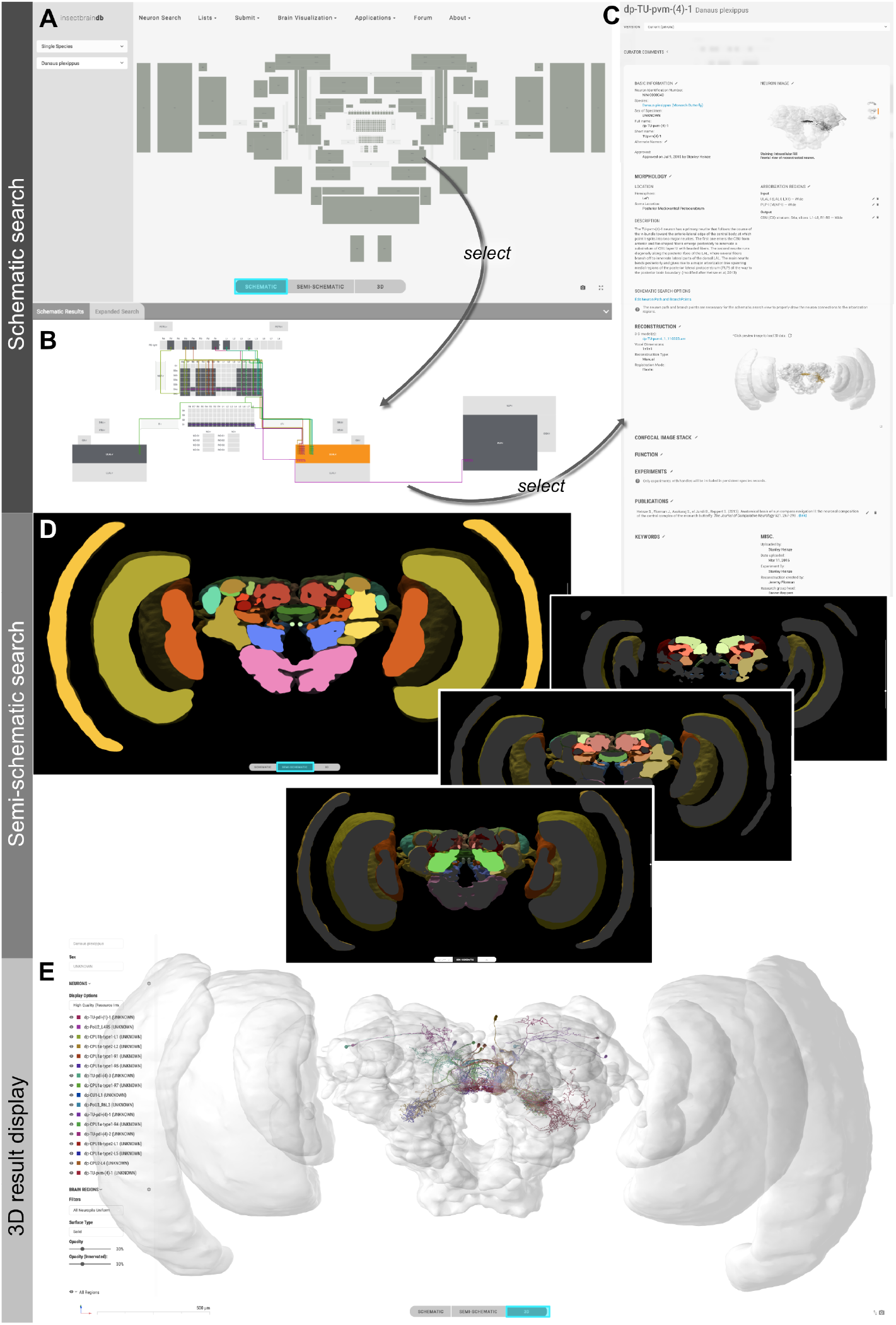
Neuron search in the *IBdb*. A. Screenshot of schematic brain search interface (Monarch butterfly) in single species mode. Selecting a neuropil will reveal all neurons connected to that neuropil. B. Schematic wiring diagram view of search result; orange neuropil was queried. Selecting an individual neuron will reveal the profile page of that cell type. C. Example of neuron profile page with anatomical information. Confocal image stack and functional data are not listed in this example. D. Semi-schematic search interface. The section view is scrollable and allows the user to query individual neuropils for connected neurons by clicking the cross section. The inserts show the results view at three levels of the brain. Neuropils connected to the queried neuropil are highlighted. Switching to the schematic view will then show the neurons as wiring diagram. Switching to the 3D mode will show registered neurons in 3D. E. The 3D results viewer allows one to view all neurons registered into a common reference frame; example from Monarch butterfly (data from ***Heinze and Reppert (2012)***). The user can continuously switch between the three modes (schematic, semi-schematic, 3D). **Figure 2-Figure supplement 1.** Schematic outline of generic brain and variations across species. **Figure 2-Figure supplement 2.** Brain structures in the database and their hierarchical organization.

If more than one species is subject to a query, a schematic brain is generated that displays the commonly shared features of the species involved. For both single and multi-species search, when selecting a specific brain region, all neurons in the *IBdb* that connect to this region become visualized by a dynamically drawn wiring diagram (***Figure 2***B). Filters can be applied to narrow down search results according to neuron polarity, functional class, etc. Individual neurons in the wiring diagram can be selected to reveal the neuron’s profile page. Here, all available information for this cell type is displayed, including links to deposited experiment entries (***Figure 2***C). The schematic display of search results can visualize any neuron in the database, only requiring that a neuron is annotated with respect to the brain regions it innervates.

For single species queries, two more modes for visualizing search results are available: the semi-schematic view and the 3D view. The semi-schematic view mode emphasizes the natural brain organization on the level of brain regions, while also serving as interface for launching search queries. It comprises a full series of automatically generated sections through a segmented 3D brain of a species. Each brain region present in that species’ 3D brain is shown as an interactive cross section that can be used to query neural connections of that region (***Figure 2***D). If a region is selected, all connected brain areas are highlighted and neurons resulting from this query can be visualized by changing to either the schematic view, the 3D view, or a list view. The advantage of the anatomically correct layout of this interface is that a brain region can be queried for a neuron, even if its name is not known to the researcher. This is particularly useful for regions with uncommon names that have only recently been introduced to the insect brain naming scheme (e.g. crepine, superior clamp, etc., see ***Ito et al.(2014)***). Launching a search for neurons in regions with unfamiliar names is made much easier when the search interface resembles the information a researcher has obtained from e.g. confocal images or physical brain slices. The semi-schematic mode of the database search function fulfills this demand and bridges the schematic wiring diagram view and the full 3D view.

The 3D view visualizes search results in an anatomically correct way and shows queried neurons in the context of a species’ reference brain (given that this information was added) (***Figure 2***E). It displays interactive surface models of that brain together with neuron skeletons obtained from the neuron-type’s profile page.

In the graphical search interface the search parameters are limited to anatomical information defining the neuron’s location in the brain (i.e. input and output areas). In contrast, an additional text-based search function (‘Expert Search’) allows users to query all information deposited on a neuron’s profile page. Individual search parameters can be logically combined to generate arbitrarily complex searches. The results are displayed as a list of neurons, which can be sent to populate a schematic wiring diagram view by a single click. Thus, this tool effectively combines complex search with the advantages of intuitive display of results.

Finally, whereas the emphasis of the database search lies on locating cell types, information on brain regions can also be found using identical interfaces. The schematic search option allows users to reveal brain region profile pages by selecting schematic neuropil representations. The same information can also be obtained by selecting brain regions in the semi-schematic neuropil search interface. Due to their heterogeneous nature, experiment entries can only be queried via the text-based search function (‘Expert Search’).

### Online applications and tools

To maximize the usefulness of the database we have implemented an integrated 3D viewer to deliver platform independent, high-quality data visualization without any additional software demands. Neuropil visibility can be independently switched on and off for each brain region, transparency can be freely adjusted, and colors of neurons can be changed (***Figure 3***A). Neurons can be shown either with diameter information or as simple backbones. The built-in screenshot function enables the user to capture any scene displayed in the 3D viewer and produces a high-resolution, publication-ready image with transparent background (***Figure 3***B,C).

**Figure 3.**
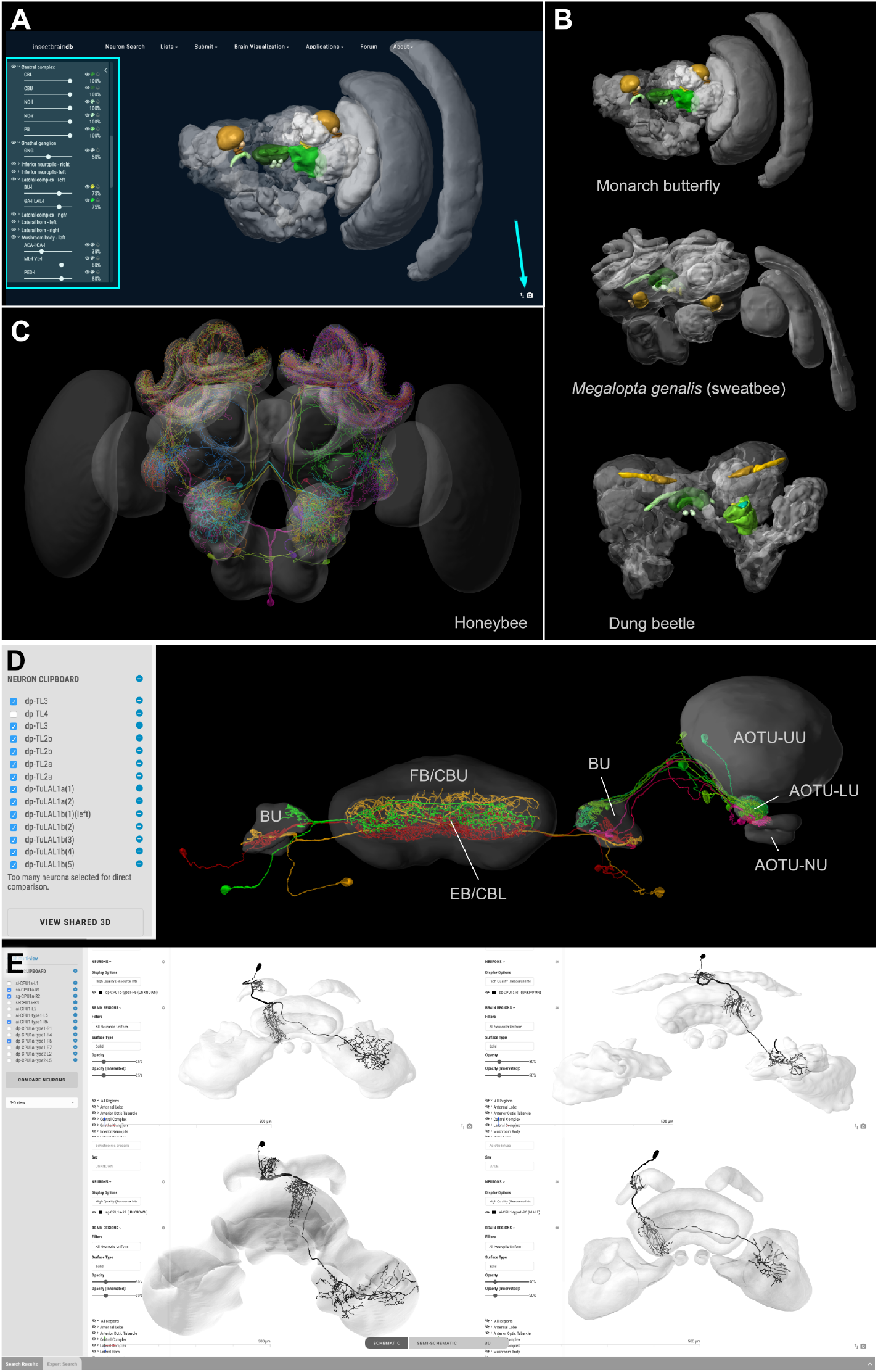
Visualization tools and applications. A. Screenshot illustrating the functionality of the 3D viewer in the insect brain database. Cyan arrow: Screenshot button. Cyan panel: Tools for adjusting appearance of neuropils. B. Examples of neuropil images generated with the *IBdb* 3D viewer, illustrating navigation relevant neuropils in three insect species (Monarch butterfly (***Heinze and Reppert, 2012***), sweat bee *Megalopta genalis* (***Stone et al., 2017***), dung beetle (***Immonen et al., 2017***). C. Neurons associated with the antennal lobe of the honeybee, generated with the *IBdb* 3D neuron viewer (data from ***Rybak (2012)***). D. Elements in the neuron clipboard (left) can be arbitrarily combined and displayed in the 3D viewer to highlight neural pathways and circuits. Shown are two parallel input pathways from the anterior optic tubercle to the ellipsoid body of the central complex in the Monarch butterfly (data from ***Heinze et al. (2013)***). E. Side-by-side neuron viewer. Screenshot showing comparison of 3D skeletons of CPU1 (PFL) neurons from four species (top left: Monarch butterfly (***Heinze et al., 2013***); bottom left: desert locust (***el Jundi et al., 2009a***); top right: Dung beetle (***el Jundi et al., 2015***); bottom right: Bogong moth (***de Vries et al., 2017***)).

The *IBdb* allows users to not only locate neuronal morphologies quickly, but also to combine arbitrary neurons from any single species into a common visualization. To achieve this, we have generated a neuron clipboard, in which individual neurons from search results can be stored temporarily (***Figure 3***D). Any subset of cells in the clipboard can be sent to the 3D viewer, as long as all neurons belong to the same species, i.e. can be displayed using the same reference brain. The desired configuration of neurons and neuropils can be generated using the interactive tools of the viewer and the screenshot function can be used to create a high-resolution image to be used for illustration purposes (e.g. reviews, conference talks, teaching).

Additionally, we have embedded a function to directly compare up to four neurons side by side on screen. Any neuron located in the neuron clipboard can be chosen to be included in this comparison. The comparison uses the 3D view, the profile image, or the confocal stack located on the respective neurons’ profile pages. This function is ideally suited to compare homologous neurons from across species to quickly assess differences and shared features of these cells. The four-window 3D viewer retains all functions ofthe normal full screen 3D viewer and thus also allows the capture of high resolution screen shots of each of the neurons being compared (***Figure 3***E).

The data in the database are suited for many applications, including more sophisticated ones. To provide direct access to all levels of the data in the *IBdb* we have created an API interface, specifying how to automatically draw data from the database via web-browser based apps. Applications produced by third parties that use this function can be embedded directly on the *IBdb* website, once they are approved by the site administrators. Applications envisioned are, for example, quantitative comparisons of both single neuron morphologies and neuropils between species, direct online multi-compartment modeling of neurons deposited in the database, or virtual reality interfaces that allow exploration of anatomical data in a 3D virtual reality environment. Over time, we hope that our unified platform will stimulate the insect neuroscience community to generate a collection of online tools to analyze and explore neuroanatomical and physiological data deposited in the *IBdb*, thereby allowing straight-forward meta-analysis of all raw data deposited in the database. As an offline tool, the Natverse package in R already offers the possibility to explore *IBdb* data (***Bates et al., 2020***).

### Contribution of data

All data on the internet is public. This also applies to any data publicly available in the *IBdb*. Driven by the requirement to obtain data persistency and implemented by the use of handles, no data can be removed from the database once it is public (and thus citable). For all data, the contributor retains ownership and holds the copyright to her/his data. The publishing is performed explicitly by the owners, not by the database administrators, and the license attached to each dataset is a Creative Commons Attribution Non Commercial 4.0 International (CC BY NC 4.0). Thus, when data in the *IBdb* are downloaded for reuse, the original work that underlies these data has to be credited together with the *IBdb* as the source. When data are used to generate images with the help of the *IBdb*, these images are licensed via Creative Commons Attribution 4.0 International (CC BY 4.0), i.e. they can be used in any publication as long as the original data owners and the *IBdb* are credited.

Data can be contributed by registered users at all levels of the database, i.e. species, brain regions, neurons, and experiments. The process is similar on all levels but requires more expert knowledge the higher in the hierarchy the data reside. In the following sections we will briefly illustrate the main principles of how to contribute data (for full instructions see the Online User Guide).

#### Species

To create a new species, a profile page has to be created, which then has to be populated with data. While photographs, bibliography, and text descriptions of the species are desirable and strongly encouraged, the most important next step is to generate a schematic brain (***Figure 4***A). This will ensure that brain structure entries are created in the database, a prerequisite for neuropil based search and 3D brain region identity. To generate a schematic brain, we have created the ‘Brain Builder’ tool on the *IBdb* website, which provides templates based on either the generic insect brain, or related species already deposited in the database. The user can simply copy an existing brain, associate it with the new species and modify it to match any unique features of the new species.

**Figure 4.**
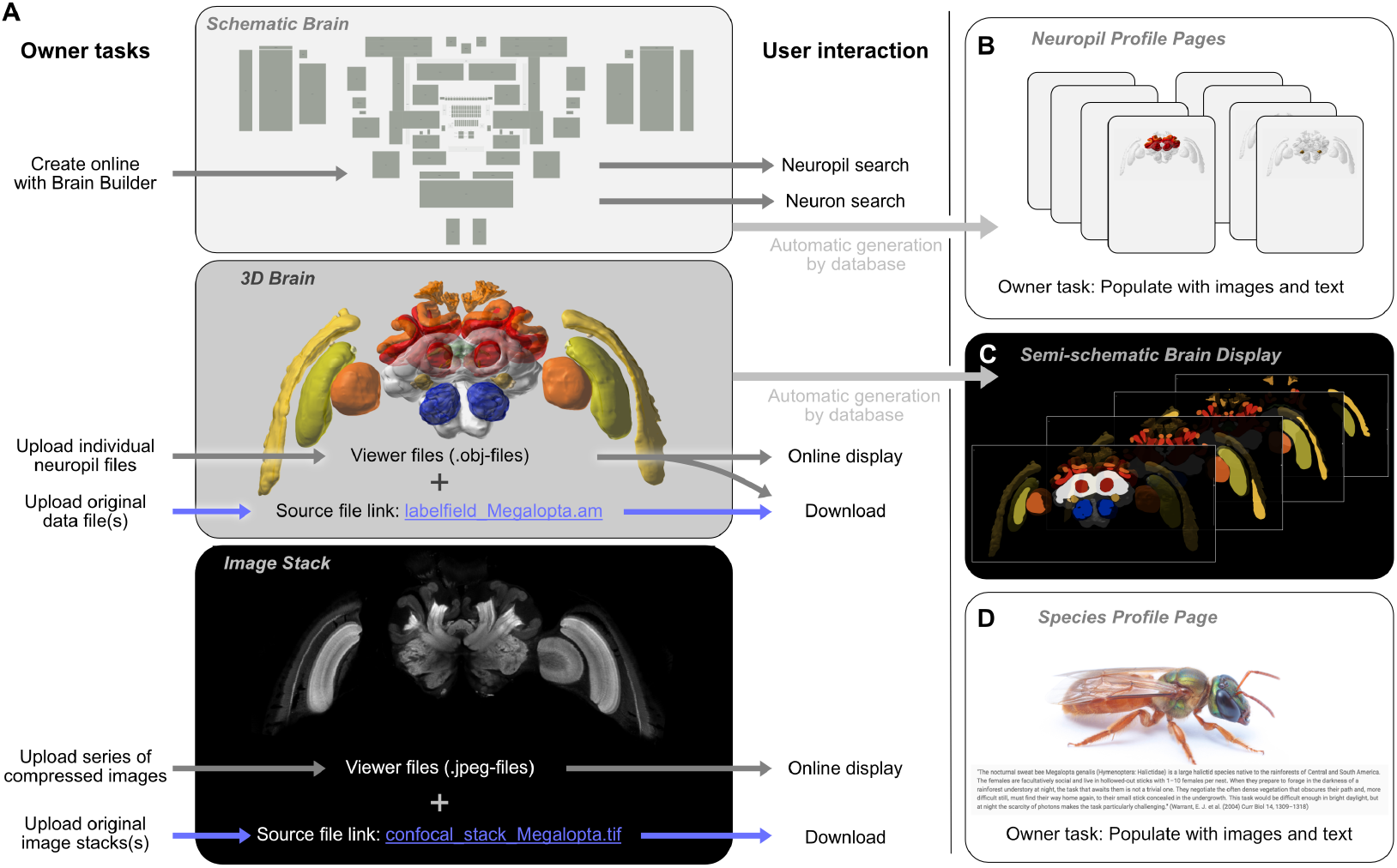
Contributing a species to the *IBdb*. A. Three main elements have to be created for each new species: the schematic brain, the 3D brain and an image stack. The schematic brain is generated directly on the *IBdb* website using the ‘Brain Builder’, while the other two elements are uploaded. For each, both source files and viewer files are needed. Viewer files are used for online display, while source files can be downloaded by users. B. Neuropil profile pages are automatically generated when creating the schematic brain. They have to be populated with images and texts by the user. C. The semi-schematic brain is automatically generated based on the provided 3D brain. D. The species profile page must be populated with images, texts and a bibliography to provide context for the species. Photograph reproduced with permission from Ajay Narendra. *Megalopta genalis* data from ***Stone et al.(2017)***

**Figure 5.**
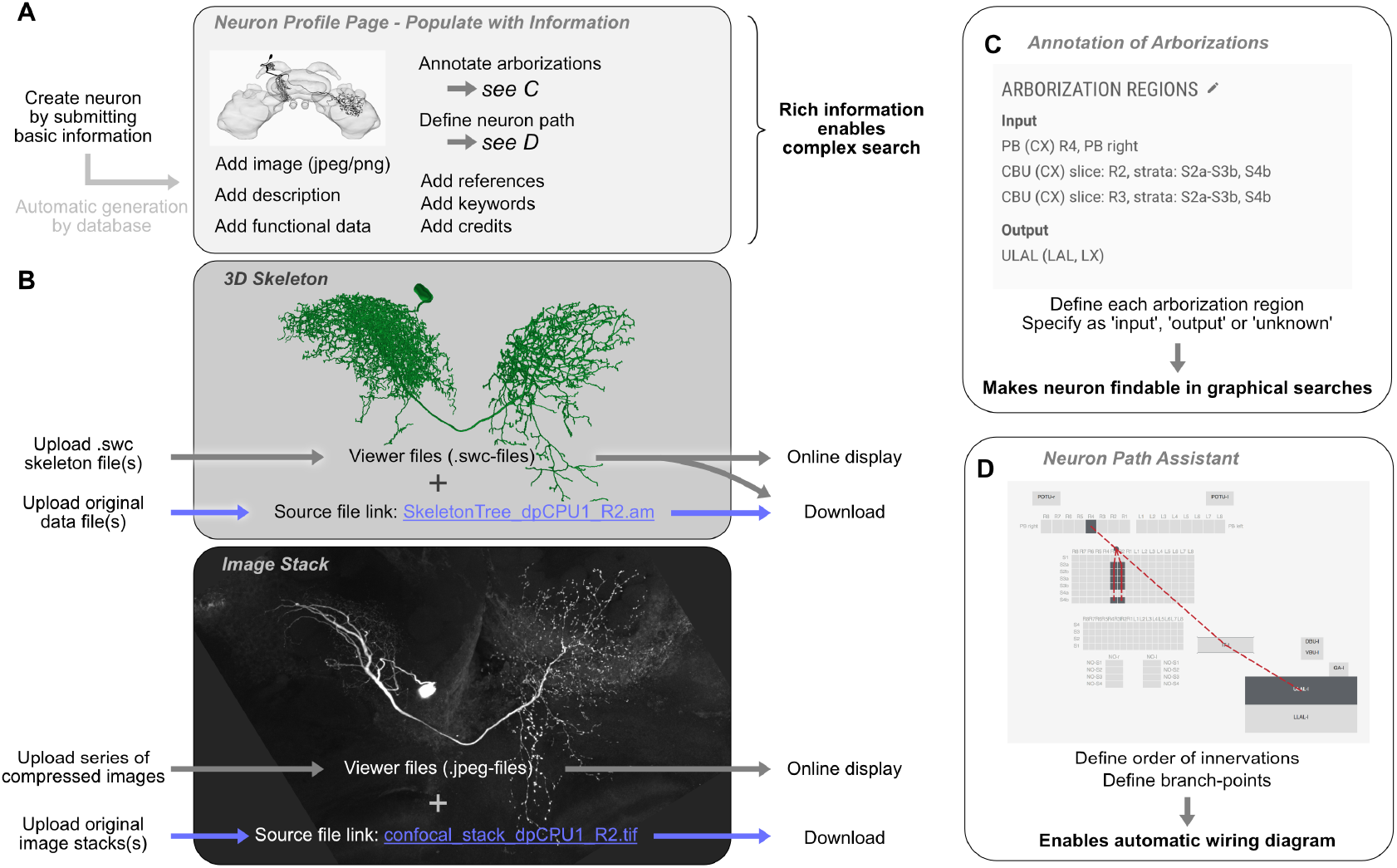
Contributing a neuron type to the *IBdb*. A. A new neuron type can be created by submitting a neuron form containing basic information. This generates a neuron profile page that then has to be populated with information by the owner. Each entry is findable by the expanded search function. B. 3D-skeletons are added as swc-files (online display) and source files (download). Confocal image stacks are uploaded as jpeg series for online display and as original data files for download. C. All arborization regions of the neuron must be defined (at the level specified in the species’ schematic brain) and labeled as either input, output or unknown polarity. D. To enable automatic drawing of wiring diagrams in schematic search results, the order of innervation of neuropils and the branch-points of the neuron must be defined using the path assistant.

Once the profile page and the schematic brain is created, a 3D brain can be uploaded to illustrate the brain organization of the new species and to serve as reference brain for neuron display (***Figure 4**A*). For all uploaded data, the database distinguishes between source files and display files. Source files contain the 3D reconstruction in a format that the researcher would like to make available for others in the field. The second set of files, the display files, are required for automatic online display and must constitute the surface models of each neuropil (.obj-format). Each brain region model is tagged with a unique neuropil identity, so that the schematic brain regions and the 3D surface models will be linked to the identical brain-structure entry. Finally, following the same principles as for the 3D brain, an image stack can be uploaded to the profile page as well (***Figure 4***A). This can be any representative dataset that illustrates the layout of the species’ brain (e.g. confocal stack, μCT image series, serial sections with any other technique).

#### Brain structures

Brain structure entries are automatically generated when defining the schematic brain search interface for a new species (***Figure 4***B). These profile pages are automatically populated by a 3D brain in which the relevant region is highlighted, a brain structure tree that reveals the relative location of the respective neuropil in the hierarchy of the species’ brain, and with links to neuron entries associated with each brain region. All remaining data have to be manually added. These are mostly descriptive in nature and encompass images, text-based descriptions, and volumetric data.

#### Neurons

Contribution of neurons follows a similar procedure as the contribution of new species (***Figure 5***).The user creates a profile page for the new cell type that subsequently has to be populated with information. To make a neuron findable in the database, its arborization regions in the brain have to be defined (***Figure 5***C). Within a graphical user interface, these regions are chosen from the brain structures available in the schematic brain of the respective species. One arborization entry has to be created for each branching domain of the neuron, leaving no part of the neuron un-annotated. To enable the automatic generation of a wiring diagram view of the new neuron for displaying schematic search results, an outline of its branching structure has to be generated in an embedded tool called the neuron-path assistant (***Figure 5***D). This branch tree defines which neuropils are innervated in which order and where main branch points are located.

The remaining procedure for neuron contribution is largely identical to species contribution and follows the dual approach towards source data and display data for 3D reconstruction and image stacks (***Figure 5***B). All other information, i.e. images, bibliography, keywords, representative functional data, transmitter content, and textual descriptions, can be added to the profile page at any time prior to publication.

Importantly, the *IBdb* can be used to house data that have been obtained by classical methods, e.g. camera lucida drawings of Golgi impregnated neurons. While no 3D information is available in those cases, drawings can be uploaded as images after which annotation of the neuron’s morphology (arborization regions) is performed as described. These neurons will therefore become findable in the schematic search interface and will be added to the publicly available pool of neuronal data.

#### Experiments

Experiment entries are created by adding them directly to the profile page to which they are linked (species, brain structure, or neuron). The automatically generated experiment profile page must then be filled with basic meta-information about the experiment (date, what was done, who did it), after which a series of files can be uploaded. These files can be in any format and are made available for download. This allows users to provide not only the raw data of any experiment, but also, for example, analysis scripts, custom made equipment-control software, and analysis results. Image files can be selected for direct online display on the experiment profile page to allow online examination of the data.

### Curation and administration

The database is managed via a group of voluntary curators, a scientific administrator, and a technical administrator. Importantly, no single person curates all data in the *IBdb*, but each species is managed by a specialized curator, who is an expert for that species. This distributed curation system ensures that no single person is responsible for too many datasets, and that no curator has to evaluate data outside their area of expertise. To additionally reduce the workload for species with many entries, more than one curator can be assigned to any given species. The scientific administrator (the lead author of this publication) oversees the curators, while technical administration is carried out by the technical administrator (last author of this publication). The technical administrator is the only person who has potential access to all data in the database.

The responsibility of the scientific administrator is to approve new species and to train and support the curators for individual species. The responsibility of each curator is to approve new neuron-type datasets and to re-evaluate major updates of these. Curator approval entails checking for formal errors in the submitted data, ensuring that new data do not accidentally duplicate already existing data, and that the quality of the data meets the standards acceptable for a particular species. To ensure swift correction of any issues, we have implemented a private communication channel between curator and data owner. This was realized through a commenting function that enables the curator to post comments on a neuron page, which are only visible to the data owner. The owner can then directly respond to the comments and any issues raised can be resolved.

To enable all users to provide feedback and to discuss topics relevant to other database users, we have added a discussion forum directly to the *IBdb* website. This forum is intended as a means for reporting potential bugs, suggesting new *IBdb* features, or for discussing scientific content (methods for data processing or acquisition, requests for literature, staining protocols etc.).

### The *IBdb* as tool for data management and data deposition

Each database entry has to be explicitly published by the contributor. In the process, it is approved by either the database administrator (species), or the species’ curator (neurons). While this procedure was initially intended only as a quality control measure to prevent incomplete or inaccurate data from compromising the database, we have developed it into a unique feature: the *IBdb* private mode. Before a dataset is made public, it is invisible to all other users, curators and the scientific database administrator. The dataset can thus be updated and even deleted. This creates the potential of using the *IBdb* to deposit data while they are being collected or prepared for publication in a research paper, i.e. for data management (***Figure 6***A).

**Figure 6.**
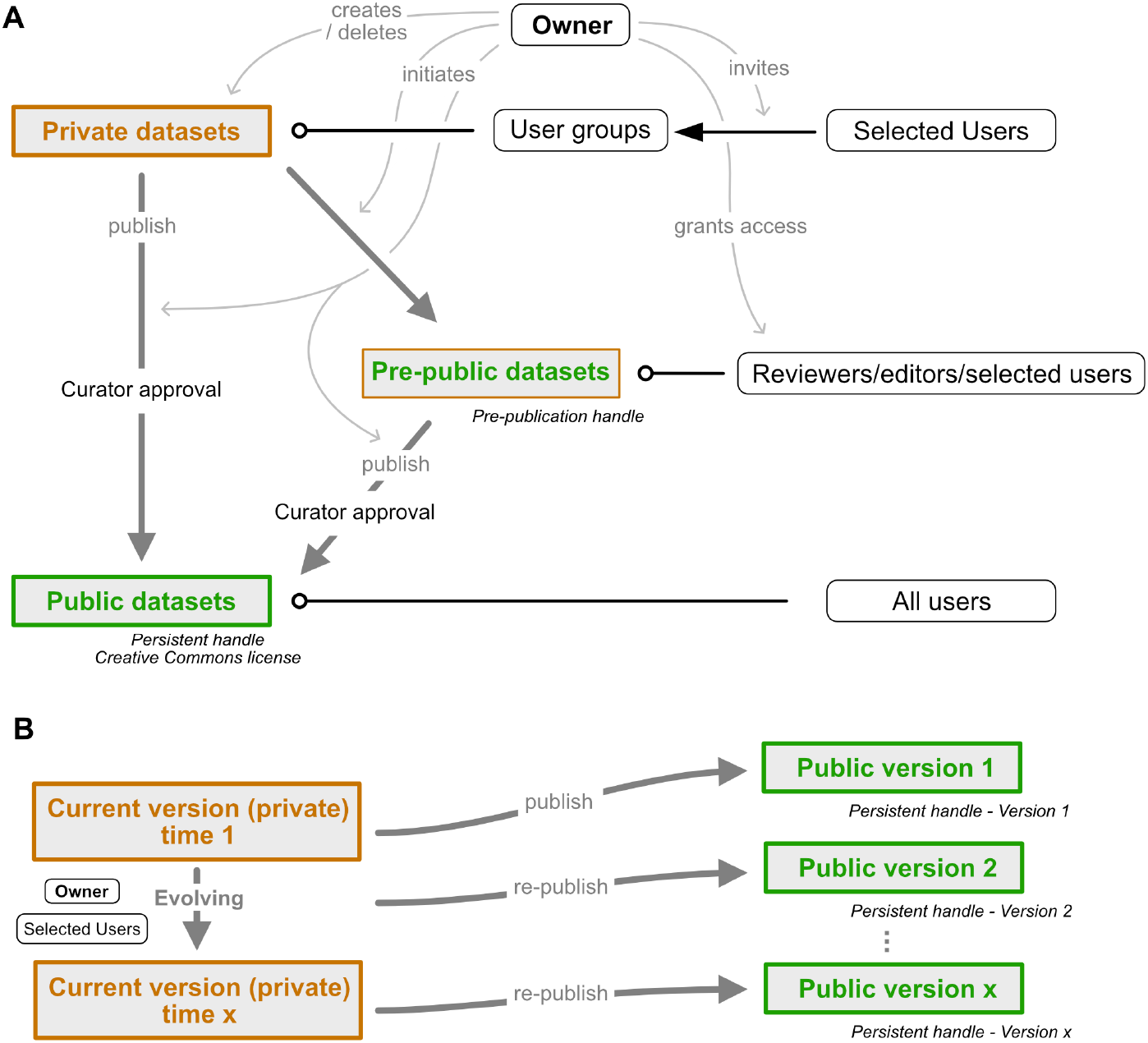
Dataset publication concept. A. Interconnection of private, public and pre-public datasets. Private datasets can be viewed and edited by members of user groups with access granted to a particular dataset. Pre-public datasets can be viewed by anyone in possession of the pre-publication handle (distributed by the data owner, e.g. within a manuscript). Public datasets can be accessed by all users. Publication of data cannot be undone as persistent handles are generated. Grey arrows indicate control actions employed by the dataset owner. B. Re-publication strategy of evolving datasets. A current version of each public dataset remains present in the owner’s private database mode and can be edited at wish. Once sufficient updates have accumulated, the dataset can be re-published. A new version of the persistent handle is assigned and the now public dataset (version 2) becomes locked. Datasets can be edited by the owner and anyone who has been granted permission to edit by the owner.

To facilitate the use of the *IBdb* as a data management tool, we have enabled three operational modes of the database site: private, public, and mixed. Any user logged in can thus choose to either access only (own) private data, only public data, or both. The first mode turns the *IBdb* into a data management site for ongoing research, the second mode is the default mode for viewing publicly available data, and the third mode allows the user to compare their own unpublished data with public data.

As efficient data management requires researchers from the same laboratory, as well as collaborators, to have access to relevant unpublished data, each user can grant access to their own private datasets (***Figure 7***). To this end, a user can create a user group and invite other database users to join. Datasets can then be added and made visible or editable to all members of the group. These data can either comprise individual entries at all levels of the database, or collections of entries defined by common features.

**Figure 7.**
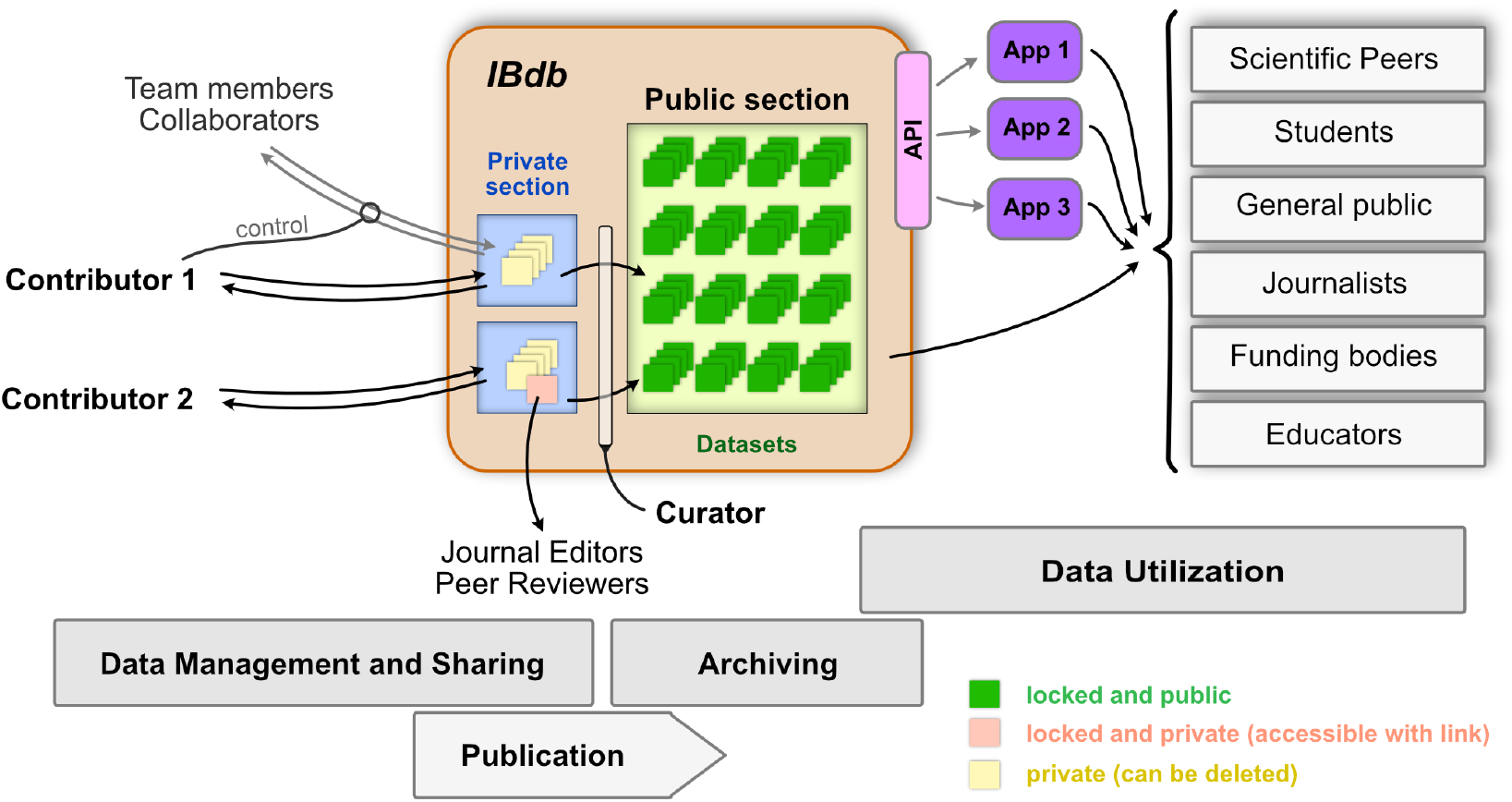
The Insect Brain Database and the possible interactions between users and deposited data. The private sections of the database are accessible to only the owner of the data, and datasets within this section can be shared with team members and collaborators. As these datasets are unlocked, they can continuously be updated and also deleted. Upon publication and curator approval, datasets become locked (persistent) and are archived in the public section of the database. As an intermediate step, datasets can be pre-published (locked but private) and made available to journal editors and peer reviewers when including datasets in manuscripts of journal articles. Data in the public section of the database is accessible directly for all interested users (relevant user groups are shown on the right). Additionally, an application programming interface (API) allows automated access of public data, which can therefore be used by third party applications (illustrated as ‘App 1-3’) for generating specific user experiences with additional capabilities, for instance in the context of teaching.

Finally, users are often reluctant to make datasets available to the public before they are included in a research paper, yet, these data should be available to editors and anonymous reviewers. We have therefore created the possibility for ‘pre-publishing’ database entries (***Figure 6***A). This function allows a user to assign a persistent handle to a dataset (e.g. a neuron profile page) without approval by the curator and without making the dataset findable through search or lists in the *IBdb*. Editors and reviewers of the research paper (anyone in possession of the handle) then have direct access to the linked pages. Once the manuscript is accepted for publication, the user can publish the respective datasets, obtain curator approval, and thus make them findable in the *IBdb* public mode.

To avoid having to separately provide numerous independent handles when sharing data, individual entries can be grouped into datasets. These receive a unique link that grants access to the entire collection. Only public or pre-public data can be included in datasets.

Public entries of the *IBdb* are maintained in a dual way; the persistent version is locked and cannot be changed, whereas a second, current version remains visible to the owner and to all members of user groups with appropriate access rights (***Figure 6***B). This current version is fully private and can be freely edited or expanded. Importantly, no data that is already part of a public version can be deleted. Rather, when for instance a confocal image stack should be replaced by a better one, the old stack can be archived, so that it will not be visible in new versions of the dataset, but will remain present in the database for display of earlier versions. Once all required updates of a dataset have been made, the edited version can be re-published and will be assigned a new version of the persistent identifier. This new version is now also locked and any further edits will again have to be done in the current version of the dataset. This ensures that all data that have been assigned a persistent identifier will remain valid and accessible, while at the same time allowing each entry to evolve. The described strategy of publishing and re-publishing and the associated duality of persistent and current versions are implemented at the levels of species, neurons and experiments.

## Discussion

### Specific problems solved by the *IBdb*

Previous and current online databases hosting insect neuroscience data have suffered from several shortcomings: Most severely for old databases, a lack of maintenance often quickly led to outdated file formats, rendering the deposited information no longer compatible for viewing with current web browsers (anticipated by ***Ito (2010)***). Second, while some databases originally allowed interactive viewing of the data, no possibility for contribution of one’s own data existed and data download was limited to very few files, e.g. ***Brandt et al. (2005) Kurylas et al. (2008)***. Usability of larger databases was generally impaired by a layout that often required expert knowledge to be able to launch meaningful database queries or to understand search results (e.g.FlyCircuit, Invertebrate Brain Platform (now:Comparative Neuroscience Platform)). This not only applies to old databases, but the restriction to individual species and often highly complex interfaces limit the potential user base to specialists even in cutting edge databases such asVFB or visualization tools such as FruitFlyBrainObservatory. Finally, the limitation to purely anatomical data, including in the major current cross-species database NeuroMorpho.org, does not account for one of the key advantages of insect neuroscience: the high level of tractable structure-function relations.

The *IBdb* addresses each of these issues. Firstly, we have developed the database software to be independent of the operating system and type of web browser used, as well as to not rely on any third party plugins. Additionally, we implemented the database as a classic, relational database without experimental data structures (e.g. intelligent, adaptive search), aiming at maximal robustness. Having created a conceptionally simple software that uses standard web-technology with standard file-formats makes continued compatibility and technical maintenance comparably simple.

Second, we have invested substantial effort in making the *IBdb* intuitive to use and visually attractive to provide a positive user experience. The latter factor should not be underestimated in its importance. One of the problems encountered in previous databases were user interfaces that were difficult to use, creating an immediate negative experience when attempting to use a site for the first time and therefore reducing the motivation to interact with it. As for commercial software, we aspired to generate a user interface that is largely self-explanatory, provides immediate visual feedback when a user action was successful, and which clearly shows what actions can be performed. Several years of beta-testing by a multitude of users have streamlined the site to a point at which interacting with the *IBdb* is both straight forward and fun.

As the success of the database depends on many users sharing their data, we aimed at making contributing data as intuitive and as easy as searching and visualizing data. We have thus simplified the data contribution process to a point where only very limited anatomical knowledge is needed, aiming at enabling physiologists without deep anatomical training to submit data as well.

Finally, to our knowledge the *IBdb* is the first database in the field of invertebrate neuroscience that combines functional data with anatomical data, and at multiple levels ranging from entire brains to single neurons. This ability, together with the possibility to deposit not only representative data but concrete sets of experiments, provides an opportunity for anatomists to directly interpret their findings in a functional context, as well as allowing physiologists to tether their findings to a coherent anatomical framework that automatically generates context for any functional data. The ease of use of the *IBdb*, combined with housing functional and anatomical data, has the potential to facilitate interactions between expert anatomists and physiologists and thereby strengthen structure-function analysis across the diversity of insect brains.

### Motivation to contribute

The landscape in the insect neuroscience research community has changed dramatically, and most relevant funding bodies are in the process of implementing open data mandates or have already done so (e.g. European Research Council, National Institutes of Health, Wellcome Trust, etc). Thus, the initial driving force for data deposition is much larger compared to that surrounding earlier database attempts. Yet, why should researchers use the *IBdb* for meeting these new mandates rather than other available databases? Different from open databases, e.g. Figshare.org, the *IBdb* is dedicated to insect neuroscience and thus provides all tools required to manage, annotate, and cross-link the specific data formats generated in this field. However, it is not rooted within a single laboratory, nor a single species, thus providing a framework for data from a broad research community. The unique possibility for cross-species comparison and the combination of anatomical and functional data additionally broadens the relevance of the *IBdb*.

We have implemented a range of tools enabling the visualization of data in fast, flexible, and effortless ways. This saves considerable time compared to other available data visualization tools, in particular for complex 3D neuron data (e.g. Amira). The data contributed are also immediately incorporated into the framework of existing data. Outside the *IBdb* these data are distributed across many publications. Comparison of one’s own data to any published data would entail contacting authors, obtaining files in unpredictable formats and finding ways to compare them to one’s own work within the software a research group is currently using. The *IBdb* solves these issues and delivers such comparisons within seconds.

Crucially, these advantages are already present immediately after data upload, prior to publication. Via the private mode of the *IBdb*, individual neurons can already be compared to their counterparts in other species while datasets are being obtained, enabling the user to generate visualizations suitable for conference contributions and publication figures. Importantly, the dual function of the IBdb as data repository and data management tool eliminates the need to reformat and prepare datasets for publication, a process that is required when submitting datasets to websites dedicated to only data deposition. The IBdb therefore provides a streamlined and integrated experience from data acquisition to publication, aimed a minimizing researcher workload.

### Scalability and long-term financial sustainability

The *IBdb* is designed for long-term accumulation of data by many contributing research groups, requiring maintenance of high technical standards and financial sustainability. The first issue is covered by an ongoing agreement with the web-developers that built and maintained the database software over the period of the last six years. Financially, voluntary contributions from research groups that initiated the database have paid for its creation. As the maintenance costs are a small fraction of the development costs, it will be easily possible to run the database within the framework of the existing service agreement for at least the next five years without any changes required. However, when the data volume increases substantially, the static costs of housing the data will increase accordingly. While keeping all public, persistent data available free of charge is mandatory (given that the *IBdb* functions as a public data repository), maintaining the *IBdb* as a free data management tool, i.e. allowing unlimited private data for each user, will likely become unsustainable over time. If this becomes a problem, free space in the private section of the database will be limited. All space required beyond a certain limit will have to be rented to directly offset the costs for maintaining and administering these data.

To anticipate the slowly growing costs of housing the database due to increasing data volume, we aim at eventually relocating the data from the currently used commercial Amazon cloud platform to an academic server that is provided at minimal costs or free of charge. To this end we have ensured that the *IBdb* does not depend on any core functions of the Amazon cloud storage service, enabling to move the database to a new location with comparably moderate effort.

### *IBdb* usage for outreach and teaching

While highly useful for classroom teaching of structure-function relations in neural systems, the *IBdb* has proven to be invaluable to introduce new members of a research team to the basic layout of brains, neurons, and neural circuits in a particular species. Using the database serves as an easy (and fun) access point to available information on a research species, including key publications, and therefore saves significant effort when writing review papers, PhD thesis introductions, background sections for travel grants, etc. While this is true for established researchers as well, it is especially true for younger scientists and students who are new to the field or are at the beginning of their careers.

Finally, the *IBdb* provides the possibility for anyone to access original research data in intuitive and attractive ways (***Figure 7***). This provides opportunities to design teaching assignments for neuroscience students to carry out meta-analyses. With access to the data in the *IBdb* via the application programming interface (API), we have provided the possibility for third parties to develop dedicated teaching tools that provide streamlined methods to use the data for specific classroom exercises. Beyond researchers and students, journalists, interested members of the public, or members of funding bodies can also view and explore neuroscience data. Ideally, this will contribute to a more transparent understanding of what the output of science is and could spark increased interest in insect neuroscience.

### *Drosophila* and Interoperability with Virtual Fly Brain

The *IBdb* does intentionally not include *Drosophila melanogaster* as a species. This is because a huge amount of effort has been spent developing highly efficient resources for this widely used model system and, as a result, the Virtual Fly Brain (VFB) resource has been created ***(Osumi-Sutherlandet al., 2014)***. It bundles data from several older *Drosophila* databases (e.g.FlyCircuit) to the most recent connectomics datasets (***Scheffer et al., 2020***). Serving as the main repository for anatomical data from the *Drosophila* brain it has become the main site to locate GAL4 driver lines, single cell morphologies, and synaptic connectivity data. It contains tens of thousands of datasets and is designed to specifically meet the needs of the *Drosophila* research community. By being less specialized, the *IBdb* has a wider scope. We are hosting many species and include both functional and anatomical data. We also do not require neuronal anatomies to be registered to a reference brain, if this is not possible for some reason. This opens the *IBdb* up to more diverse data, but as a result cannot provide most of the specialized services that VFB can deliver (e.g. automatic bridging registrations of 3D data between different reference brains). Importantly, both databases have converged on a highly similar ontological framework. As the brain nomenclature used by the *IBdb* and VFB is identical, neuropil identities can be mapped across both databases and direct links can be provided between them. Ideally, this will eventually enable a user to launch a query in the *IBdb* and directly link the results to the corresponding data inVFB (and vice versa). This interoperability will make maximal use of both complementary resources, without duplicating functionality.

### Widening the scope towards other animal groups

The framework we have generated with the *IBdb* is not limited to housing insect brain data. Without major modifications it would be equally suited for hosting data from other animal taxa. While the intuitive, schematic search engine would not be useful for comparing species that do not share a common basic brain outline (i.e. a relevant ‘generic brain’), the text-based expanded search could allow the construction of queries across multiple groups, e.g. searches according to functional terms. We are currently conceptualizing the expansion of the database towards including spiders and envision that crustaceans and other arthropods would be logical next groups.

The *IBdb* therefore provides not only a tool for the insect neuroscience community to facilitate data management, data visualization, transparency of results and effective teaching, but it can easily be expanded towards related fields. Additionally, it might also serve as a blueprint for how to set up similar databases in unrelated research areas. In principle, the strategies used in the *IBdb* are applicable to any scientific field that can be linked to a hierarchical, ontological framework.

## Methods and Materials

### Data location

All web infrastructure is hosted by Amazon web services on servers located in Frankfurt, Germany. Data is stored using a PostgreSQL relational database hosted by the Amazon Relational Database Service and files are stored using the Amazon S3 object storage service. The servers hosting the website and the local HANDLE system are running in Amazon EC2 containers, which runs Linux. Resources communicate using Amazon Virtual Private Cloud.

### Database framework

The database structure and interaction is managed by a python based Django application. User authentication, permissions and data security are also managed within the Django application. A NGINX web server hosts static content and serves as a reverse proxy for dynamic content served by a uWSGI application server hosting the project’s Django application. Asynchronous tasks are implemented using the Celery distributed task queue and RabbitMQ message broker.

Data is externally accessible via a web API delivering content in the JSON format to the front-end web application. The web API was implemented with Django and the Django REST framework.

Longterm data persistency is provided to allow users to reference information or profile pages on the site in scientific publications and other external media in a static state, while continuing to allow data to be updated as more information is acquired. When a request is made by a user for a persistent copy of a dataset to be created, a copy of the data related to the current state of the dataset is serialized and parsed into JSON. A persistent unique identifier is then assigned (HANDLE). The JSON data, HANDLE and additional metadata is recorded in a separate table and can no longer be modified. All files associated with the persistent dataset are marked as locked in the database and can no longer be modified by the user. The recorded state of that dataset can be accessed and viewed on the site using the url associated with the assigned HANDLE. The original data copied to create the persistent dataset can be modified without effecting the persistent dataset. Additional files may also be added, but will not be reflected in earlier persistent records.

### Graphical user interface

The front-end of the database is primarily implemented using the Angular web framework in Type-script, HTML and SASS (CSS extension language). The Typescript, HTML and SASS are compiled and bundled with the Angular CLI using WebPack to create the distributed application files targeting ECMAScript 2015 capable browsers. Graphical consistency is targeted for browsers using Webkit and Gecko based layout engines adhering to web standards.

The web based three-dimensional viewer was implemented using Typescript, WebGL and the Three.js three dimensional graphics framework. The two-dimensional schematic view, brain designer and path designer was implemented using Typescript, the Canvas API and Paper.js vector graphics scripting framework.

### Security measures

User to server communication is protected by the Hyper Text Transfer Protocol Secure (HTTPS). User authentication is managed by the Django authentication system. Access to data is restricted by object based permissions limited to authorized users through the web interface or API arbitrated by the Django server.

File downloads of protected content stored on Amazon S3 are accessed using time limited urls, assigned to an authorized user at the time of a download request by the Django server. Files are directly downloaded from the S3 storage to the user using the temporary URL. The process is seamless to the end user who need only be logged into the website and click the link associate with the intended file. Downloads are logged and accessible to the owner of the data being accessed. Files can be uploaded to the S3 object storage only by authorized users with a time limited URL provided by the server. The upload is logged and associated with the contributing user. Publicly available thumbnail images and other reduced quality images are stored separately in a publicly accessible (read-only) S3 bucket.

Content explicitly designated as public is accessible through the graphical user interface or via the API to any authorized visitor. Private content is only accessible to authorized users given permission to access the data.

The PostgreSQL database is protected by a firewall allowing access only via the Amazon Virtual Private Cloud and is not open to direct access via the web.

### Nomenclature and Brain Hierarchy

All names for brain areas are in line with previous research. The brain regions of the generic insect brain follow the new insect brain nomenclature introduced for *Drosophila* by***Ito et al. (2014)***. Accordingly, we have established three hierarchical levels of brain regions: super-regions, neuropils, and sub-regions. Super-regions are stereotypical and can be expected to comprise the ground pattern of the brain in all insects (although some might be reduced in certain species). The only exception to the ***Ito et al. (2014)*** scheme is the anterior optic tubercle, which we have raised to the level of super-region, given its prominence and distinct nature in most insect species. Sub-regions are often specific to individual species and therefore, if such regions were defined, we used the names given to them within the relevant species. We did not unify e.g. names of the mushroom body calyx divisions across species, as this would firstly imply homology where there might not be any, and second, novel naming schemes will have to be developed by the community and not be imposed by a data repository. Anticipating that changes to brain names can and will happen in the future, all names, as well as the level of a region within a hierarchy, can be modified.

Within some neuropils, regular, repeating elements can be found, usually defined as columns and layers. We have implemented such a system in the central complex, i.e. without having to define an array of sub-regions, several strata and orthogonal slices (following the new brain nomenclature) can be generated. The default number of slices in the generic brain is 16, assuming that this number is the ancestral state of this region.

Neuron names follow the conventions within each species, as there is no common naming scheme for insect brain neurons yet in place. However, we provide the possibility to define several alternative names for each cell type to allow the parallel use of names. This is possible as the identity of a neuron is linked to the persistent ID, and not to the neuron’s name. Given that we house neurons from multiple species, we add a prefix to the full name of each cell type specifying the species, e.g. ‘am’ for *Apis mellifera*.

## Acknowledgments

We are indebted to the many test users of the *IBdb* who patiently located bugs and inconsistencies and thereby helped to streamline the database outline and make the user interface more intuitive. We also would like to thank all members of the Heinze lab for many helpful discussions that improved the *IBdb* and this manuscript. Funding was provided from the following sources: Air Force Office of Scientific Research (Grant No. FA9550-14-1-0242) (to E.W and S.H.), European Research Council (ERC) under the European Union’s Horizon 2020 research and innovation program (to S.H., grant agreement no. 714599 and M.D., grant agreement no. 817535), the German Research Foundation (EL784/1-1 to B.elJ.; HO 950/24-1, HO 950/25-1 and HO 950/26-1 to U.H.; Me365/34 to R.M.), startup grant from the University of Würzburg to K.P., Norwegian Research Council (project number 287052) to B.B., Freie Universität Berlin and Zukunftskolleg University Konstanz (supporting R.M.), and the Swedish Research Council (project number 2014 - 04623) to M. D..

---

**Figure 1-Figure supplement 1.**
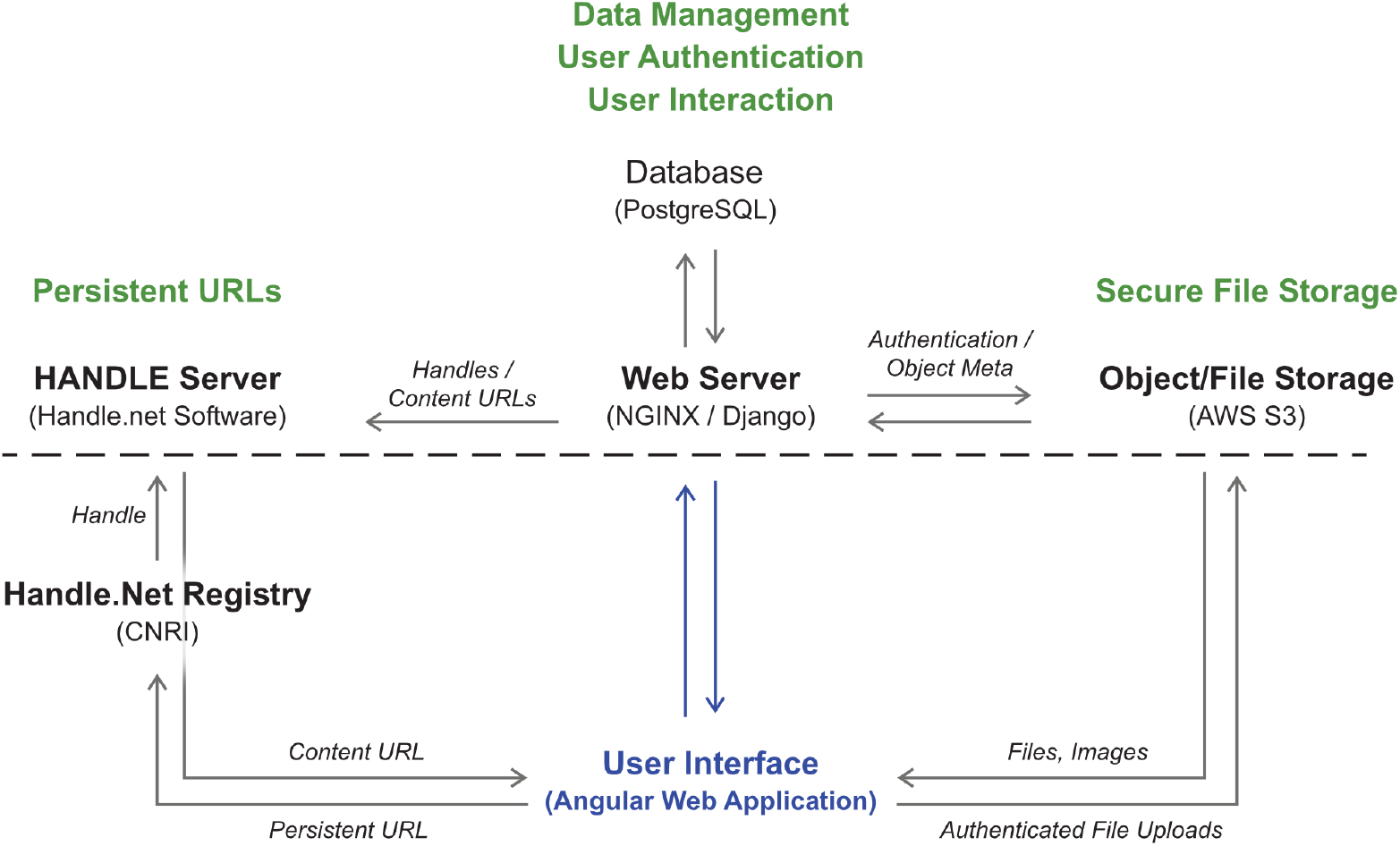
Illustration of the interactions between the user interface, the web server, database, data storage and handles management.

**Figure 2-Figure supplement 1.**
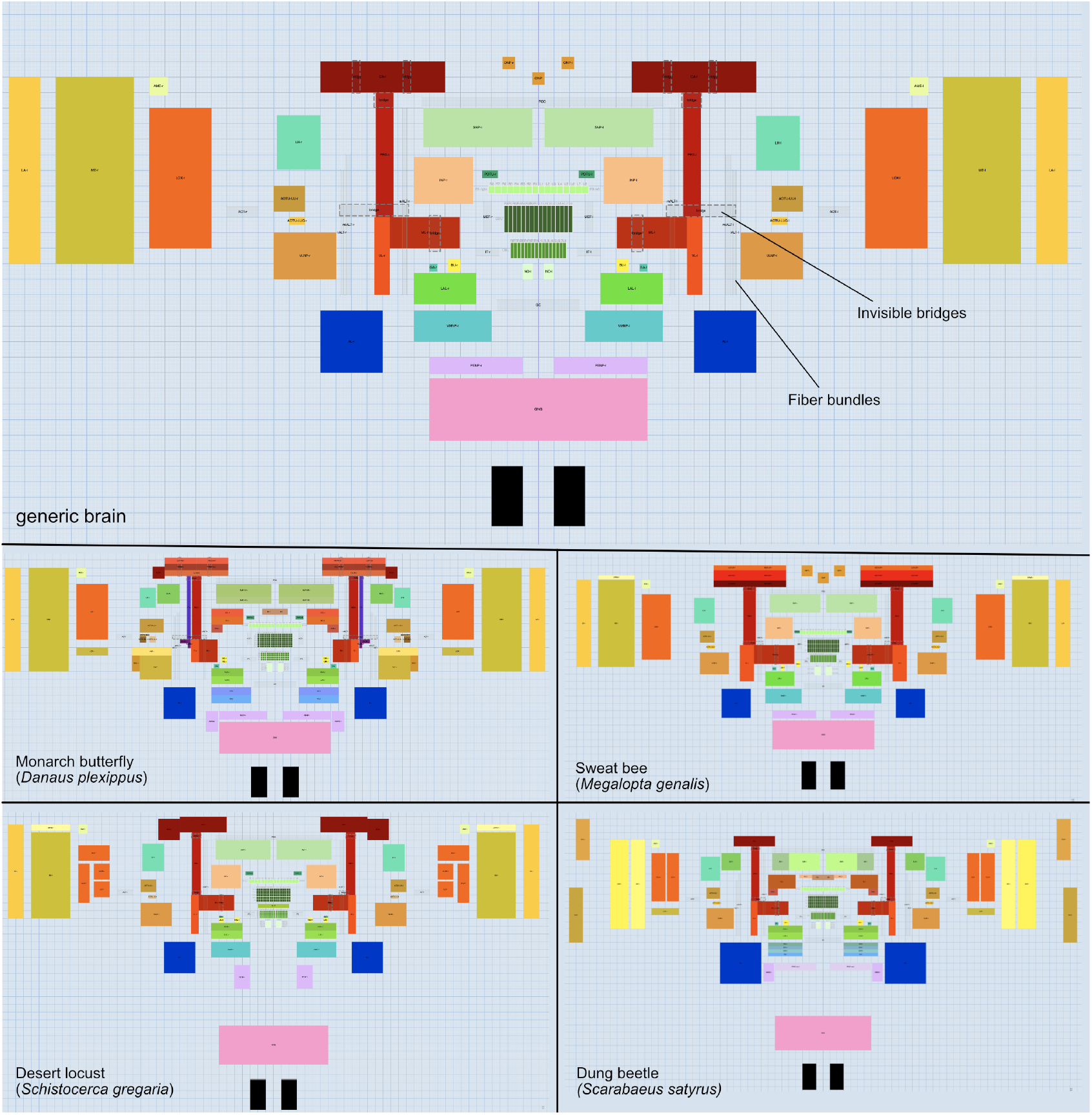
The brain builder module enables to generate the schematic brain models required for the graphical search functions. It allows the user to draw brain regions (tagged as either super-region, neuropil, or sub-region) as well as fiber bundles and invisible bridges. The latter provide shortcuts for neurons in the wiring diagram view (schematic search results), preventing overly convoluted neuron paths. The top panel shows the generic brain that is used as the template for all other species and which combines brain regions and fiber bundles shared by the majority of insect species. The bottom four panels illustrate the modifications of this template needed to account for species-specific features of four insect species examples.

**Figure 2-Figure supplement 2.**
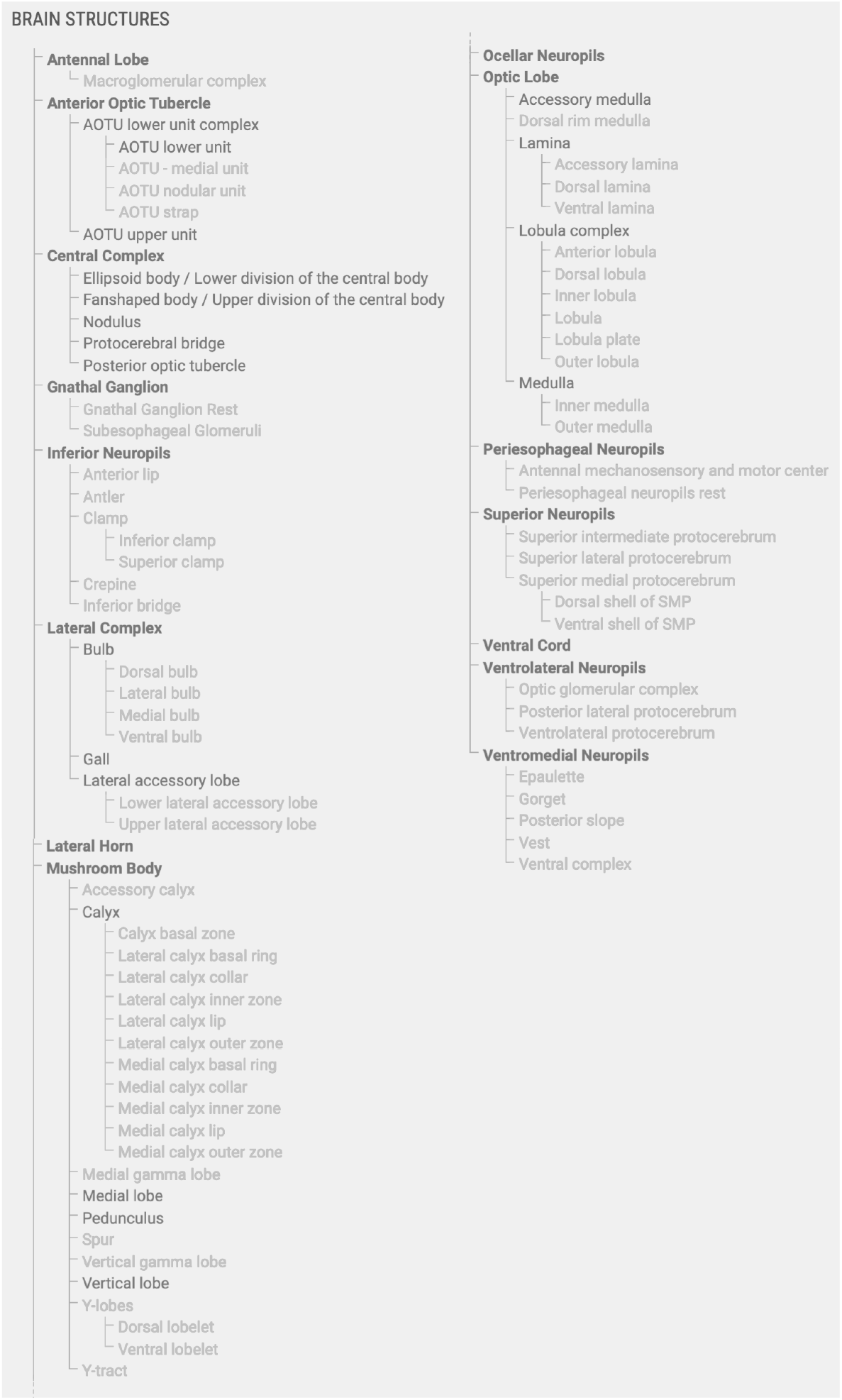
Brain structures included in the insect brain database. All regions shown in black font are included in the generic insect brain that serves as a template for all other species and as a fall-back option for cross-species search. Bold regions are super-regions, which consist of neuropils, which in turn can be divided into sub-regions. Note that neuropils and sub-regions can be species-specific (grey font) and no single species contains all listed structures.

## References

Armstrong JD, Kaiser K, Müller A, Fischbach KF, Merchant N, Strausfeld NJ. Flybrain, an on-line atlas and database of the Drosophila nervous system. Neuron. 1995 Jul; 15(1):17–20.

Ascoli GA, Donohue DE, Halavi M. NeuroMorpho.Org: a central resource for neuronal morphologies. The Journal of Neuroscience. 2007 Aug; 27(35):9247–9251.

Bates AS, Manton JD, Jagannathan SR, Costa M, Schlegel P, Rohlfing T, Jefferis GS. The natverse, a versatile toolbox for combining and analysing neuroanatomical data. eLife. 2020 Apr; 9.

Brandt R, Rohlfing T, Rybak J, Krofczik S, Maye A, Westerhoff M, Hege HC, Menzel R. Three-dimensional average-shape atlas of the honeybee brain and its applications. The Journal of Comparative Neurology. 2005 Nov; 492(1):1–19.

Chiang AS, Lin CY, Chuang CC, Chang HM, Hsieh CH, Yeh CW, Shih CT, Wu JJ, Wang GT, Chen YC, Wu CC, Chen GY, Ching YT, Lee PC, Lin HH, Wu CC, Hsu HW, Huang YA, Chen JY, Chiang HJ, et al. Three-dimensional re-construction of brain-wide wiring networks in Drosophila at single-cell resolution. Current Biology. 2011 Jan; 21(1):1–11.

Dreyer D, Vitt H, Dippel S, Goetz B, el Jundi B, Kollmann M, Huetteroth W, Schachtner J. 3D Standard Brain of the Red Flour Beetle Tribolium Castaneum: A Tool to Study Metamorphic Development and Adult Plasticity. Frontiers in Systems Neuroscience. 2010; 4:3.

Heinze S, Florman J, Asokaraj S, el Jundi B, Reppert SM. Anatomical basis of sun compass navigation II: the neuronal composition of the central complex of the monarch butterfly. The Journal of Comparative Neurology. 2013 Feb; 521(2):267–298.

Heinze S, Reppert SM. Anatomical basis of sun compass navigation I: The general layout of the monarch butterfly brain. The Journal of Comparative Neurology. 2012 Jun; 520(8):1599–1628.

Immonen EV, Dacke M, Heinze S, el Jundi B. Anatomical organization of the brain of a diurnal and a nocturnal dung beetle. Journal of Comparative Neurology. 2017 Jun; 525(8):1879–1908.

Ito K. Technical and organizational considerations for the long-term maintenance and development of digital brain atlases and web-based databases. Frontiers in Systems Neuroscience. 2010 Jan; 4:26.

Ito K, Shinomiya K, Ito M, Armstrong JD, Boyan G, Hartenstein V, Harzsch S, Heisenberg M, Homberg U, Jenett A, Keshishian H, Restifo LL, Rössler W, Simpson JH, Strausfeld NJ, Strauss R, Vosshall LB, Insect Brain Name Working Group. A systematic nomenclature for the insect brain. Neuron. 2014 Feb; 81(4):755–765.

el Jundi B, Heinze S, Lenschow C, Kurylas A, Rohlfing T, Homberg U. The Locust Standard Brain: A 3D Standard of the Central Complex as a Platform for Neural Network Analysis. Frontiers in Systems Neuroscience. 2009; 3:21.

el Jundi B, Huetteroth W, Kurylas AE, Schachtner J. Anisometric brain dimorphism revisited: Implementation of a volumetric 3D standard brain in Manduca sexta. The Journal of Comparative Neurology. 2009 Nov; 517(2):210–225.

el Jundi B, Warrant EJ, Byrne MJ, Khaldy L, Baird E, Smolka J, Dacke M. Neural coding underlying the cue preference for celestial orientation. Proceedings of the National Academy of Sciences of the United States of America. 2015 Sep; 112(36):11395–11400.

Kurylas AE, Rohlfing T, Krofczik S, Jenett A, Homberg U. Standardized atlas of the brain of the desert locust, Schistocerca gregaria. Cell and Tissue Research. 2008 Jul; 333(1):125–145.

Mayernik MS. Open data: Accountability and transparency. Big Data and Society. 2017 jul; 4:1–5.

Osumi-Sutherland D, Costa M, Court R, O’Kane CJ. Virtual Fly Brain - Using OWL to support the mapping and genetic dissection of the Drosophila brain. CEUR workshop proceedings. 2014 Oct; 1265:85–96.

Rybak J. The Digital Honey Bee Brain Atlas. In: Honeybee Neurobiology and Behavior Springer Netherlands; 2012.p. 125–140.

Scheffer LK, Xu CS, Januszewski M, Lu Z, Takemura SY, Hayworth KJ, Huang GB, Shinomiya K, Maitlin-Shepard J, Berg S, Clements J, Hubbard PM, Katz WT, Umayam L, Zhao T, Ackerman D, Blakely T, Bogovic J, Dolafi T, Kainmueller D, et al. A connectome and analysis of the adult Drosophila central brain. eLife. 2020 Sep; 9:1073–83.

Stone T, Webb B, Adden A, Weddig NB, Honkanen A, Templin R, Wcislo W, Scimeca L, Warrant EJ, Heinze S. An anatomically constrained model for path integration in the bee brain. Current Biology. 2017 Oct; 27(20):3069–3085.e11.

de Vries L, Pfeiffer K, Trebels B, Adden AK, Green K, Warrant EJ, Heinze S. Comparison of Navigation-Related Brain Regions in Migratory versus Non-Migratory Noctuid Moths. Frontiers in Behavioral Neuroscience. 2017; 11:158.

Wilkinson MD, Dumontier M, Aalbersberg IJJ, Appleton G, Axton M, Baak A, Blomberg N, Boiten JW, da Silva Santos LB, Bourne PE, Bouwman J, Brookes AJ, Clark T, Crosas M, Dillo I, Dumon O, Edmunds S, Evelo CT, Finkers R, Gonzalez-Beltran A, et al. The FAIR Guiding Principles for scientific data management and stewardship. Scientific data. 2016 Mar; 3(1):160018–9.

